# Health behaviors among male and female university students in Cambodia: a cross-sectional survey

**DOI:** 10.1101/742718

**Authors:** Say Sok, Khuondyla Pal, Sovannary Tuot, Rosa Yi, Pheak Chhoun, Siyan Yi

**Author notes:** Co-authors’ emails: SS, KP ST RY, PC.

## Abstract

**Background:** Students go through a transition when they enter university, which involves major individual and contextual changes in every domain of life that may lead to several behavioral and health problems. This paper describes a wide range of health behaviors among male and female university students in Cambodia.

**Methods:** A cross-sectional survey was conducted in 2015 among students randomly selected from the Royal University of Phnom Penh and University of Battambang. Health-related behaviors in different domains were collected using a structured questionnaire. Chi-square test, Fisher’s exact test, or independent Student’s *t*-test was used as appropriate to describe and compare the variables among male and female students.

**Results:** This study included 1359 students, of whom 50.8% were male, and the mean age was 21.3 years (SD= 2.3). Of the total, 79.5% reported not having any vigorous-intensity activities, 25.9% not having moderate-intensity activities, and 33.5% not having walked continuously for 10 minutes during the past seven days. The prevalence of substance use was low with 38.3% currently drinking alcohol, 1.1% smoking tobacco, and 0.4% using an illicit drug during the past 12 months. About one in ten (10.6%) reported having sexual intercourse, with a mean number of partners of 2.1 (SD= 2.4) during the past 12 months, and 42.4% not using a condom in the last intercourse. Only 7.1% reported having been diagnosed with a sexually transmitted infection in the past 12 months; of whom, 60% sought for treatment. About one-third (33.6%) reported eating fast food once or twice, and 5.3% having it three times or more over the last week. More than half (55.6%) had one to two servings of fruits and vegetables daily, and 9.9% did not eat any fruits and vegetables over the last week.

**Conclusions:** We found that the prevalence of sexual risk behaviors and substance use was plausibly low among university students in this study. However, the rates of inactive lifestyle and unhealthy food consumption were concerning. Public policy and universities should promote healthy behaviors among the students. The interventions may take advantage of and expend upon the good health behaviors and consider gender differences.

## Introduction

Globally, chronic diseases such as cancer, stroke, heart disease, and diabetes are the major causes of death [1]. Behavioral risk factors including tobacco smoking, alcohol consumption, physical inactivity, sedentary behaviors, and obesity are major determinants of adult chronic diseases, morbidity, and mortality [1–4]. In addition to the disease burden attributed to single chronic behavioral risk factors, a growing body of evidence suggests that many behavioral risk factors co-occur among youths, including university students [5–7] and that their combinations yield greater risks for chronic diseases than the sum of their individual independent effects [7–9].

People living in developing countries are mostly affected by health risks and behaviors associated with poverty such as under-nutrition and unsafe sex [10]. With economic growth and increase in life expectancies, major risks to health shift from these risks to other more contemporary risks, e.g. obesity and physical inactivity, other diet-related factors such as fast food and sugary product consumption, and tobacco and alcohol-related risks [11]. Besides coping with the heavy burden of diseases associated with underdevelopment, these countries thus need to fight with the growing burden of non-communicable diseases as well [10].

Different health risk behaviors have been shown to be associated with several important factors, including economic growth, mobility, and low self-esteem. Studies have shown that economic growth can lead to a rise in obesity or over nutrition, which suggests that university students aiming for higher economic mobility may be at risk for obesity [12]. This population needs to be aware of the myriads of health risks for diseases in order to prevent further incidence [6,13]. There are also many predictors for suicidal behaviors, alcohol and other drug problems which include psychiatric disorders, substance abuse, lack of social support, negative family environment, major life stressors, peer pressure and conformity, and demographic factors [14,15].

Health behaviors formed during childhood and adolescence, including during university, can have a significant impact on the occurrence of future illnesses and may continue into adulthood and beyond [16,17]. While the full etiology of any of these diseases has yet to be understood, behavioral factors such as tobacco use, exercise, diet, alcohol consumption, and preventive health checks are strongly implicated as risk factors [3,18,19].

Like the trend in general youths, health risk behaviors among students are more inclined towards the negative too. Research into the health behaviors of young European university students indicates a growing trend towards a less healthy lifestyle [6,7,16]. Heavy drinking and other risky behaviors, for example, tend to increase during this academic transition [20,21]. Many studies from around the world have identified different health behaviors among university students such as transport safety, violence, smoking, the use of alcohol and illegal drugs, sexual behaviors, unhealthy eating, weight control, and the lack of practice of physical exercise [6,7,16,22–24]. Like anywhere else, students in Cambodia go through the same transition when they enter university, which involves major individual and contextual change in every domain of life. However, there is no known study that examines health risk behaviors among Cambodian university students. A few studies on risky sexual behaviors, alcohol consumption, and tobacco use among other Cambodian youth groups, however, indicates that some health risk behaviors are rather a concern [25,26].

Studies of health behaviors, especially of university students, in low- and middle-income countries are important since the mortality and morbidity from common causes such as coronary heart disease and cancers and adoption of healthy lifestyles differ widely across countries [6,7,10,27], and not much is known about the situation in Cambodia. The National Institute for Health and Care Excellence, the United Kingdom, stressed that social and economic conditions can prevent people from changing their behaviors to improve their health and can also reinforce behaviors that damage it [28]. Besides the great disparity of social and economic conditions within the country, being one of the fastest growing economies would intensify the impact of social and economic conditions on lifestyle and behaviors of young populations in Cambodia.

The rationale for studying university students is that this population is faced with a lot of transitions in their academic and living environments [7], which might lead to either positive or negative behaviors. They need to adapt to new changes, brought about by the academic transition, and have more freedom over their health and lifestyles. This important transition provides a good opportunity to adopt healthy behaviors or otherwise [29]. Research has shown that many university students throughout the world engage in health risk behaviors, and such can have long-term implications for their health and lifestyle [6,16,17,30]. Studies regarding health behaviors related to non-communicable diseases in developing countries are still scarce [7]. Further, this population subgroup will occupy important positions in the government and society, including ones that promote public health. It is therefore important to understand their health behaviors and to promote remedial actions [7].

The aim of this study is to assess a wide range of health behaviors among university students in Cambodia. Broadly speaking, the article focuses on six areas of basic health behaviors: physical activities, substance use, sexual behaviors, eating behaviors, sleeping behaviors, and driving behaviors. Variables covered in this study are presented in the sub-section on variables and measurements in more details below.

## Materials and Methods

### Ethics statement

The National Ethics Committee for Health Research of the Ministry of Health, Cambodia, approved the study protocol and materials (No. 191NECHR). Participation in this study was voluntary. In the process of obtaining their written informed consent, participants were made clear that they could refuse or discontinue their participation at any time and for any reason. The confidentiality and privacy of the respondents were protected by administering the questionnaires in a private premise and by excluding personal identifiers in the survey.

### Study design and settings

This cross-sectional study was conducted in June and July 2015 among a sample of undergraduate students randomly selected from the Royal University of Phnom Penh and University of Battambang. These universities were purposively selected to partake in this research. The Royal University of Phnom Penh is the oldest and largest university in the country, and the University of Battambang is one of the most established universities in northwestern Cambodia.

### Participants and sample size

Epi Info (Atlanta, GA, United States) was used to calculate the sample size. There were approximately 168000 students registered in the higher education institutions [31]. Because the prevalence of health risk behaviors among university students in Cambodia was not known, a 50% rate was used for the calculation to prevent any underestimated prevalence. Based on a 95% confidence level (CI) and a 5% margin of error, the minimum required sample size was 767 university students. Adjusted for 10% of incomplete response, missing data and rejection rate, the final minimum required sample size was 850.

### Sampling and data collection procedures

Undergraduate students from first to fourth years from all departments in the two universities were eligible and selected to participate in the study. A multi-stage cluster sampling method was used to select the participants. First, two universities were conveniently selected for administration and logistics purposes. All departments in the selected universities were included in the study. In each department, a proportional to size sampling method was used to select the students to meet the required sample size. Participants were randomly selected from a name list of students in each department, and a personal identification number was assigned to each selected student. On the designated date of data collection, all selected students were approached by our trained data collectors, and questionnaires with an information sheet were delivered to them. Students were asked for a written informed consent. The self-administered questionnaire was developed in English, then translated into Khmer, and back translated into English by local experts. The questionnaire was pre-tested as part of data collection training with 10 male and 10 female students at the Royal University of Phnom Penh, who were later excluded from the main study.

### Variables and measurements

The design of the core part of the self-reported questionnaire was informed by the Health Behavior Survey [32], the National College Health Risk Behavior Survey (the United States) [33], and the Global School-based Student Health Survey [34]. There were five sections in the survey. The first section focused on socio-demographic characteristics of the participants including age (continuous), gender (male, female), marital status (single, married, cohabiting), year of study in the university, living situation (with parent, relative, friend, partner, or alone), perceived family economic status (wealthy, quite well-off, not very well-off, quite poor), academic performance (excellent, good, fair, poor), weight (in kilograms, continuous), and height (in centimeters, continuous). The second section collected information on vigorous-intensity and moderate physical activities during the last seven days using the International Physical Activity Questionnaire short version (IPAQ-S7S) [35].

Substance use, including tobacco use, alcohol drinking, and drug use, was covered in the third section. Variables included current use of tobacco (yes, no), experience of daily tobacco use (yes, no), frequency of smoking (not smoking, once or twice per week, weekly), current status of alcohol drinking (non-drinker, very occasional drinker, occasional drinker, regular drinker), average number of alcohol drinks during the past two weeks (cans or bottles for beer and glasses for wine, continuous), and the use of illicit drugs (yes, no) during past 12 months.

Sexual behaviors were the focus of the fourth section. The participants were asked questions on whether they had sexual intercourse during the past 12 months (yes, no), age at first sexual intercourse (continuous), number of sexual partners during the past 12 months (continuous), alcohol and drug use during the last sex (yes, no), condom use during the last sex (yes, no), sex in exchange for money or gifts during the past 12 months (yes, no), condom use during last sex in exchange for money or gifts, diagnosis of sexually transmitted infections (STIs) during the past 12 months (yes, no), treatment seeking behaviors for the most recent STI, and history of unwanted pregnancy in lifetime.

Eating, sleeping, and driving behaviors were covered in the last section. Overall, the variables in this section collected information on the frequency and amount of fast food, soft drinks, soda or sweet tea, high-fat snacks, margarine, butter, meat fat, fruits or vegetables, and lean protein consumed during the past week. We also asked about the average sleeping hours in 24 hours during the past week (continuous).

### Data analyses

Data were cleaned and entered into a Microsoft Excel (version 2010) database and analyzed using SPSS version 22.0 (IBM Corporation, New York, USA). Descriptive statistics were used to compute means and standard deviations for continuous variables as well as frequencies and percentages for categorical variables. Exploratory univariate and bivariate analyses were conducted to measure the distribution of frequencies of variables. Bivariate analyses were conducted using χ^2^ test (or Fisher’s exact test when a sample size was smaller than five in one cell) for categorical variables and independent Student’s *t*-test for continuous variables to compare socio-demographic characteristics, health status, and health risk behaviors by student sex.

## Results

Of 1462 students approached, 1359 completed the questionnaire (a response rate of 92.9%). The majority (98.6%) of those who declined the participation reported time constraint as the main reason. About half (50.8%) of the respondents were male, and the mean age was 21.3 years (SD= 2.3). The majority (97.9%) of the respondents were single. Of the total, 36.5% was in year one, and 70.8% reported fair academic performance. Almost half (43.8%) were living with parents, and 67.3% reported coming from a not very well-off or quite poor family. There were significant differences between male and female respondents in terms of age, living situation, family economic conditions, height, and weight (Table 1).

**Table 1.** Socio-demographic characteristics of male and female university students.

| Variables | Frequency |  |  | *p*-value* |
| --- | --- | --- | --- | --- |
|  | Total<br>*n* (%) | Male<br>*n* (%) | Female<br>*n* (%) |  |
| Age (in years, mean ± SD) | 21.3 ± 2.3 | 21.7 ± 2.5 | 21 ± 2.2 | <0.001 |
| Marital status |  |  |  | 0.88 |
| Single | 1330 (97.9) | 674 (97.7) | 656 (98.1) |  |
| Married | 27 (2.0) | 15 (2.2) | 12 (1.8) |  |
| Cohabiting | 2 (0.1) | 1 (0.1) | 1 (0.1) |  |
| Year of study in the university |  |  |  | 0.15 |
| Year 1 | 496 (36.5) | 245 (35.5) | 251 (37.5) |  |
| Year 2 | 281 (20.7) | 130 (18.8) | 151 (22.6) |  |
| Year 3 | 251 (18.5) | 137 (19.86) | 114 (17.0) |  |
| Year 4 | 331 (24.3) | 178 (25.8) | 153 (22.9) |  |
| Living arrangement |  |  |  | <0.001 |
| Parent | 595 (43.8) | 271 (39.3) | 324 (48.4) |  |
| Relative | 335 (24.6) | 173 (25.1) | 162 (24.2) |  |
| Friends | 346 (25.5) | 188 (27.2) | 158 (23.6) |  |
| Couple | 21 (1.5) | 14 (2.0) | 7 (1.0) |  |
| Alone | 18 (2.7) | 44 (6.4) | 18 (2.7) |  |
| Perceived family economic status |  |  |  | <0.001 |
| Wealthy | 1 (0.1) | 1 (0.2) | 0 (0.0) |  |
| Quite well-off | 443 (32.6) | 194 (28.2) | 249 (37.2) |  |
| Not very well off | 846 (62.3) | 449 (65.2) | 397 (59.3) |  |
| Quite poor | 68 (5.0) | 45 (6.5) | 23 (3.4) |  |
| Perceived academic performance |  |  |  | 0.92 |
| Excellent | 57 (4.2) | 29 (4.2) | 28 (4.2) |  |
| Good | 303 (22.3) | 157 (22.8) | 146 (21.8) |  |
| Faire | 962 (70.8) | 487 (70.6) | 475 (71.0) |  |
| Poor | 37 (2.7) | 17 (2.5) | 20 (3.0) |  |
| Weight (in kg, mean ± SD) | 53.0 ± 9.1 | 57.7 ± 8.5 | 48.1 ± 6.9 | <0.001 |
| Height (in cm, mean ± SD) | 163.1 ± 14.8 | 168.5 ± 18.5 | 157.5 ± 5.3 | <0.001 |

Abbreviations: n, number; SD, standard deviation.
\*Chi-square test (or Fisher's exact test when a sample size was smaller than five in one cell) was used for categorical variables and independent Student's t-test for continuous variables.

As shown in Table 2, 79.5% of the respondents had no vigorous intensity activity; 25.9% had no moderate-intensity activities; and 33.5% did not walk continuously for 10 minutes during the past seven days. More than half of those who had vigorous-intensity activities (51.1%) and those who had moderate-intensity activities (59.6%) did it in a period of less than 30 minutes. Of those who had walked continuously for at least 10 minutes, 73.1% did it in a period of less than 30 minutes. There is a statistical difference between male and female in the frequency and time spent on vigorous-intensity activities.

**Table 2.** Physical activities among male and university students in the study.

| Variables | Frequency |  |  | *p*-value |
| --- | --- | --- | --- | --- |
|  | Total<br>*n* (%) | Male<br>*n* (%) | Female<br>*n* (%) |  |
| Work involved vigorous-intensity activities for ≥10 min continuously |  |  |  | 0.001 |
| Never | 973 (71.7) | 444 (64.4) | 529 (79.5) |  |
| Had 1-3 days/week | 261 (19.3) | 159 (23.1) | 102 (15.3) |  |
| Had 4-7 days/week | 120 (8.9) | 86 (12.5) | 34 (5.1) |  |
| Time usually spent doing vigorous-intensity activities on one of those days |  |  |  | 0.001 |
| ≤30 minutes | 161 (51.1) | 89 (42.8) | 72 (67.3) |  |
| 31-60 minutes | 65 (20.6) | 53 (25.5) | 12 (11.2) |  |
| >60 minutes | 89 (28.3) | 66 (31.7) | 23 (21.5) |  |
| Work involved moderate-intensity activities for ≥10 min continuously |  |  |  | 0.61 |
| Never | 352 (25.9) | 185 (26.9) | 167 (25.0) |  |
| Had 1-3 days/week | 564 (41.6) | 278 (40.4) | 286 (42.8) |  |
| Had 4-7 days/week | 441 (32.5) | 226 (32.8) | 215 (32.2) |  |
| Time spent doing moderate-intensity activities on one of those days |  |  |  | 0.94 |
| ≤30 minutes | 514 (59.6) | 264 (60.0) | 250 (59.1) |  |
| 31-60 minutes | 204 (23.6) | 104 (23.6) | 100 (23.6) |  |
| >60 minutes | 145 (16.8) | 72 (16.4) | 73 (17.3) |  |
| Days walking for ≥10 min continuously to get to and from places |  |  |  | 0.08 |
| Never | 470 (34.7) | 247 (35.9) | 223 (33.5) |  |
| Had 1-3 days/week | 338 (24.9) | 154 (22.4) | 184 (27.6) |  |
| Had 4-7 days/week | 547 (40.4) | 288 (41.8) | 259 (38.9) |  |
| Time usually spent walking on one of those days |  |  |  | 0.19 |
| ≤30 minutes | 545 (73.1) | 261 (70.2) | 284 (75.9) |  |
| 31-60 minutes | 111 (14.9) | 60 (16.1) | 51 (13.6) |  |
| >60 minutes | 90 (12.1) | 51 (13.7) | 39 (10.4) |  |

*Abbreviations: n, number.*
*\*Chi-square test (or Fisher's exact test when a sample size was smaller than five in one cell) was used.*

Table 3 shows that 38.3% of the respondents reported currently drinking alcohol, 1.1% smoking tobacco, and 0.4% using illicit drugs during the past 12 months. Of those who reported using illicit drugs during the past 12 months, 50% used methamphetamines, 25% used heroine, and 25% used other types of drugs. Among those who reported alcohol drinking, average number of days they drank during the past two weeks was 1.3 (SD= 2.0), and average number of alcoholic drinks used during the past two weeks was 1.8 (SD= 2.9). There were statistically significant gender differences in frequency of smoking, frequency in alcohol drinking, and average number of alcoholic drinks used during the past two weeks.

**Table 3.** Substance use among male and female university students in the study.

| Variables | Frequency |  |  | *p*-Value |
| --- | --- | --- | --- | --- |
|  | Total<br>*n* (%) | Male<br>*n* (%) | Female<br>*n* (%) |  |
| Current tobacco smokers |  |  |  | 0.47 |
| Yes | 15 (1.1) | 9 (1.3) | 6 (0.9) |  |
| No | 1344 (98.9) | 681 (98.7) | 663 (99.1) |  |
| Experience of daily smoking for at least 30 days |  |  |  | 0.16 |
| Yes | 8 (0.6) | 6 (0.9) | 2 (0.3) |  |
| No | 1351 (99.4) | 684 (99.1) | 667 (99.7) |  |
| Frequency of smoking during the past 3 months |  |  |  | 0.002 |
| I don't smoke | 1343 (98.8) | 676 (98.0) | 667 (99.8) |  |
| Once or twice | 13 (1.0) | 12 (1.7) | 1 (0.1) |  |
| Weekly | 3 (0.2) | 2 (0.3) | 1 (0.1) |  |
| Would you describe yourself as: |  |  |  | <0.001 |
| Non-drinker | 838 (61.7) | 274 (39.7) | 564 (84.3) |  |
| Very occasional drinker | 367 (27.0) | 292 (42.3) | 75 (11.2) |  |
| Occasional drinker | 151 (11.1) | 121 (17.5) | 30 (4.5) |  |
| Regular drinker | 3 (0.2) | 3 (0.4) | 0 (0.0) |  |
| Average number of days drinking alcohol during the past 2 weeks (mean ± SD) | 1.2 ± 2.0 | 1.4 ± 2.2 | 0.7 ± 2.0 | 0.001 |
| Average number of alcohol drinks<sup>†</sup> during the past 2 weeks (mean ±SD) | 1.8 ± 3.0 | 2.1 ± 3.1 | 1.1 ± 2.2 | 0.002 |
| Used illicit drugs during the past 12 months |  |  |  | 0.33 |
| Yes | 4 (0.3) | 3 (0.4) | 1 (0.1) |  |
| No | 1355 (99.7) | 687 (99.6) | 668 (99.9) |  |

Abbreviations: n, number; SD, standard deviation.
\*Chi-square test (or Fisher's exact test when a sample size was smaller than five in one cell) was used for categorical variables and independent Student's t-test for continuous variables.
<sup>†</sup>Can was a unit used for beer and glass for wine.

Regarding sexual behaviors (Table 4), 10.6% of the respondents reported having sexual intercourse during the past 12 months with an average number of sexual partners of (SD= 2.4). Among those who responded, 42.4% reported not using condom, and 6.3% reported using alcohol during their last sexual intercourse. Only 9.2% reported having sex in exchange for money or gifts. Less than one in 10 (7.1%) reported having been diagnosed with an STI in the past 12 months; of whom, 60% sought for treatment for the most recent STI. Among students who reported having sexual intercourse in the past 12 months, 12.7% reported having been or made someone pregnant in lifetime. There were significant gender differences in sexual encounter, mean number of sexual partners, condom use, and experience in having sex with commercial partners.

**Table 4.** Sexual behaviors among male and female university students in the study.

| Variables | Frequency |  |  | p-Value |
| --- | --- | --- | --- | --- |
|  | Total<br>n (%) | Male<br>n (%) | Female<br>n (%) |  |
| Had sexual intercourse in the past 12 months |  |  |  | <0.001 |
| Yes | 144 (10.6) | 119 (17.3) | 25 (3.7) |  |
| No | 1215 (89.4) | 571 (82.7) | 644 (96.3) |  |
| Alcohol use during last sex |  |  |  | 0.60 |
| Yes | 9 (6.3) | 8 (6.8) | 1 (4.0) |  |
| No | 134 (93.7) | 110 (93.2) | 24 (96.0) |  |
| Condom use during last sex |  |  |  | 0.004 |
| Yes | 83 (57.6) | 75 (63.0) | 8 (32.0) |  |
| No | 61 (42.4) | 44 (37.0) | 17 (68.0) |  |
| Sex in exchange for money or gifts |  |  |  | 0.08 |
| Yes | 13 (9.2) | 13 (11.2) | 0 (0) |  |
| No | 128 (90.8) | 103 (88.8) | 25 (100) |  |
| Condom use during last sex in exchange for money or gifts |  |  |  | N/A |
| Yes | 8 (61.5) | 8 (61.5) | N/A |  |
| No | 5 (38.5) | 5 (38.5) | N/A |  |
| Diagnosed with a sexually transmitted infection in the past 12 months |  |  |  | 0.51 |
| Yes | 10 (7.1) | 9 (7.8) | 1 (4.0) |  |
| No | 131 (92.9) | 107 (92.2) | 24 (96.0) |  |
| Sought for treatment for the most recent sexually transmitted infection |  |  |  | 0.39 |
| Yes | 6 (60.0) | 5 (55.6) | 1 (100) |  |
| No | 4 (40.0) | 4 (44.4) | 0 (0) |  |
| Ever made someone pregnant/been pregnant |  |  |  | 0.58 |
| Yes | 18 (12.7) | 14 (12.0) | 4 (16.0) |  |
| No | 124 (87.3) | 103 (88.0) | 21 (84.0) |  |
| Age at first sexual intercourse (in years, mean± SD) | 20.6 ± 3.2 | 20.7 ± 2.9 | 20.2 ± 4.4 | 0.47 |
| Number of partner during the past 12 months (mean± SD) | 2.1 ± 2.4 | 2.3 ± 2.6 | 1.0 ± 0.2 | 0.02 |
| Age first made someone pregnant/became pregnant (in years, mean± SD) | 23.1 ± 3.2 | 23.3 ± 3.4 | 22.3 ± 2.3 | 0.65 |

Abbreviations: n, number; SD, standard deviation.
\*Chi-square test (or Fisher's exact test when a sample size was smaller than five in one cell) was used for categorical variables and independent Student's t-test for continuous variables.

Table 5 presents eating behaviors among the participants. Of the total, 33.6% reported having fast food once or twice per week, and 5.3% having it three times or more per week. More than half (57.3%) reported drinking soft or sweet drinks once or twice per week, and 26.0% drinking three times or more per week. About two-thirds (63.9%) reported consuming high-fat snacks once or twice per week, and 13.5% consuming three times or more per week. Similarly, 51.2% reported consuming fat and its associated products daily, and 7.2% consuming a lot of them. About one in ten (9.9%) reported not consuming any fruits and vegetables, and 12.1% not consuming any lean meat in the past week.

**Table 5.** Eating, sleeping, and driving behaviors among male and female university students.

| Variables | Frequency |  |  | *p*-Value |
| --- | --- | --- | --- | --- |
|  | Total<br>*n* (%) | Male<br>*n* (%) | Female<br>*n* (%) |  |
| Times do eating fast food per week |  |  |  | <0.001 |
| 0 times | 881 (61.1) | 457 (66.2) | 374 (55.9) |  |
| 1-2 times | 456 (33.6) | 201 (29.1) | 255 (38.1) |  |
| 3 or more times | 72 (5.3) | 32 (4.6) | 40 (6.0) |  |
| Number of soft drinks, glasses of soda or sweet tea consumed per day |  |  |  | 0.01 |
| None | 226 (16.6) | 100 (14.5) | 126 (18.8) |  |
| 1-2 | 779 (57.3) | 389 (56.4) | 390 (58.3) |  |
| 3 or more | 354 (26.0) | 201 (29.1) | 153 (22.9) |  |
| Times consuming high-fat snacks per week |  |  |  | 0.001 |
| 0 times | 307 (22.6) | 174 (25.2) | 133 (19.9) |  |
| 1-2 times | 869 (63.9) | 445 (64.5) | 424 (63.4) |  |
| 3 or more times | 183 (13.5) | 71 (10.3) | 112 (16.7) |  |
| Amount of margarine, butter, meat fat consumed |  |  |  | 0.67 |
| None or very little | 565 (41.6) | 285 (41.3) | 280 (41.9) |  |
| Some | 696 (51.2) | 351 (50.9) | 345 (51.6) |  |
| A lot | 98 (7.2) | 54 (7.8) | 44 (6.6) |  |
| Number of servings of fruits or vegetables consumed per day |  |  |  | 0.006 |
| 0 servings | 134 (9.9) | 84 (12.2) | 50 (7.5) |  |
| 1-2 servings | 755 (55.6) | 385 (55.8) | 370 (55.3) |  |
| 3 or more servings | 470 (34.6) | 221 (32.0) | 249 (37.2) |  |
| Times consuming lean protein per week |  |  |  | 0.81 |
| 0 times | 164 (12.1) | 83 (12.0) | 81 (12.1) |  |
| 1-2 times | 874 (64.3) | 439 (63.6) | 435 (65.0) |  |
| 3 or more times | 321 (21.6) | 168 (24.3) | 153 (22.9) |  |
| Self-perception on body weight |  |  |  | <0.001 |
| Overweight | 408 (30.0) | 157 (22.8) | 251 (37.5) |  |
| About right | 496 (36.5) | 251 (36.4) | 245 (36.6) |  |
| Underweight | 455 (33.5) | 282 (40.8) | 173 (25.8) |  |
| Problem with sleeping in the last 30 days |  |  |  | 0.98 |
| None | 215 (15.8) | 110 (15.9) | 105 (15.7) |  |
| Mild | 454 (33.4) | 231 (33.5) | 223 (33.3) |  |
| Moderate | 587 (43.2) | 295 (42.8) | 292 (43.6) |  |
| Severe | 103 (7.6) | 54 (7.8) | 49 (7.3) |  |
| Used seatbelt when driving a car or other motor vehicle in the past 30 days |  |  |  | 0.04 |
| I did not drive | 756 (55.6) | 397 (57.5) | 359 (53.7) |  |
| Never | 211 (15.5) | 88 (12.8) | 123 (18.4) |  |
| Rarely | 93 (6.8) | 44 (6.4) | 49 (7.3) |  |
| Sometimes | 108 (7.9) | 60 (8.7) | 48 (7.2) |  |
| Most of the time | 68 (5.0) | 41 (5.9) | 27 (4.0) |  |
| Always | 123 (9.1) | 60 (8.7) | 63 (9.4) |  |
| Used a helmet when riding a bicycle in the past 30 days |  |  |  | 0.04 |
| I did not ride | 156 (11.5) | 64 (9.3) | 92 (13.8) |  |
| Never | 49 (3.6) | 22 (3.2) | 27 (4.0) |  |
| Rarely | 33 (2.4) | 19 (2.8) | 14 (2.1) |  |
| Sometimes | 51 (3.8) | 29 (4.2) | 22 (3.3) |  |
| Most of the time | 319 (23.5) | 179 (25.9) | 140 (25.9) |  |
| Always | 751 (55.3) | 377 (54.6) | 374 (55.9) |  |
| Drove a car or other motor vehicle after drinking alcohol in the past 30 days |  |  |  | <0.001 |
| I did not drive | 371 (27.3) | 151 (21.9) | 220 (32.9) |  |
| 0 times | 797 (58.6) | 370 (53.6) | 427 (63.8) |  |
| 1 time | 120 (8.8) | 105 (15.2) | 15 (2.2) |  |
| 2-3 times | 52 (3.8) | 48 (7.0) | 4 (0.6) |  |
| ≥4 times | 19 (1.4) | 16 (2.3) | 3 (0.4) |  |
| Average sleeping hours/24h during the past week (± SD) | 7.2 ± 1.5 | 7.2 ± 1.5 | 7.3 ± 1.5 | 0.15 |

Abbreviations: n, number; SD, standard deviation.
\*Chi-square test (or Fisher's exact test when a sample size was smaller than five in one cell) was used for categorical variables and independent Student's t-test for continuous variables.

As also shown in Table 5, on average, participants reported sleeping 7.2 hours per day (SD= 1.5), while 33.4% reported having mild sleeping problems, and 50.8% having moderate or severe sleeping problems. Of those who used a car in the past 12 months, 22.3% never or rarely used a seatbelt. Of those who used a motorbike, 6.0% never or rarely used a helmet. Regarding drunk driving, 14.0% reported driving or riding a vehicle after drinking at least once during the past 30 days. There were significant sex differences in the frequency of consumption of fast food and high-fat snacks, amount of soft or sweet drinks, fat and fat-associated products, and fruits or vegetables. Perception on body weight, seatbelt and helmet use, and drunk driving practice also differed significantly among the two sexes.

## Discussion

This study is among the few investigations into health behaviors of Cambodian university students. There were a comparable number of male and female respondents, and they came from all years of the undergraduate programs. While many major findings above are perhaps more or less in line with findings elsewhere or in other studies of Cambodian youths, there are some exceptions too.

A positive point about the students’ health behaviors is that the majority of the respondents were not involved in any of the major health-risk behaviors, including tobacco use, drug use, alcohol consumption, and sexual activities, and hence they could veer from negative outcomes from such behaviors. Some of these findings seem to contradict with or different from those of previous studies conducted on university students or youths, more broadly, in Cambodia [36] or elsewhere [37]. Reproductive health issues and casual sex, for instance, were reportedly higher among other Cambodian youth groups [25,38]. The self-reported consumption of alcohol seemed to be relatively lower than that of adolescents in Cambodia and in some other countries in the region as reported in other studies [6,39,40]. This may also be applied to the use of tobacco [41].

Such positive results are good for individual development and public health, given the grave consequences from their consumption or involvement on the individuals’ health and finance and on the government and society at large [11,37]. Alcohol and drug abuse, for instance, are known to be negatively associated with individuals’ and family wellbeing and can incur high societal and economic cost to the government [42], while drug use and trafficking is a barrier to achieving international development, including the Sustainable Development Goals [42]. A systematic review of alcohol consumption and disease burden found that alcohol impacts many disease outcomes (both chronic and acute) and injuries, including death [43]. Another systematic review indicated that heavy alcohol consumption in the late adolescence continues into adulthood and is associated with alcohol problems [44].

Besides, there were some positive eating behaviors among the students too according to their self-report, i.e. low consumption of fast food, soft and sweet drinks, high-fat snacks, and fat and fat-associated products, as well as quite adequate sleep. The findings on healthy dietary behaviors from other studies are inconclusive, with some indicating that students elsewhere have ‘satisfactory’ eating habits, while others reported concerning behaviors [17,45]. Public health policies in Cambodia should take advantage of and sustain these positive behaviors and try to promote such behaviors beyond their university life.

While students’ limited involvement in major health-risk behaviors and some positive health behaviors are plausible, many other negative health behaviors were concerning. These issues include relatively high consumption of soft and sweet drinks; low consumption of fruits, vegetables, and lean protein; sleep disorders; low frequency in usage of a seatbelt and helmet; and drunk driving practices. Many of these behaviors can have negative implications for individual health, as many of them are more susceptible to the modern health risks [8,9,11,16,45], which may result in big financial burden and social costs on the individuals and government in the future.

Another noticeable negative instance was the inactive lifestyles of many participants. The majority of them never had any vigorous- or moderate-intensity activities, and never walked continuously for 10 minutes or more. For those who had vigorous-intensity activities, moderate-intensity activities, and walked continuously for 10 minutes or so, the majority reported to have done so for less than 30 minutes, the level of daily moderate-intensity activity recommended by the World Health Organization for a healthy lifestyle [46]. Other studies also found that university students had inactive lifestyles [7,45]. Apart from the proactive response from the national policy, therefore, Cambodian universities may need to promote healthy lifestyles and habits; for example, through extracurricular activities and social engagement.

Differences between the two sexes can be observed in some areas. First, female students were observed to have better dietary behaviors and nutritional knowledge, perhaps due to the need to watch over one’s weight and stronger beliefs in healthy eating [17,45]. Second, there was a significant difference between male and female students in smoking frequency, frequency and amount of alcohol consumption [6], sexual activities including number of partners and condom use, frequency of fast food and high-fat snack consumption, and amount of soft and sweet drinks. Third, there is a statistically significant difference between the two sexes in the involvement of vigorous-intensity activities and time spent on the activities. Gender differences can also be seen in the amount of fruits and vegetables consumed, perception of body weight, frequency in usage of seatbelts and helmets, and frequency of drunk driving practices. Other studies also found gender differences in physical activities (male being more active) and body weight [7,45]. Given the sex differences in many outcome variables (including negative health risk behaviors), a gender lens may be considered in any potential interventions and policy considerations.

Overall, the negative health behaviors and lifestyles could present multiple health risks, especially the threat from chronic diseases and/or severe health conditions, to the participants in the future [2,3,47], depending on the intensity of their current health behaviors, risks and lifestyles, and the changes in behaviors and lifestyles in the future. Coupled with the health risks associated with economic underdevelopment, they may be facing, that is, according to the family status of a majority of the respondents, e.g., 67.3% of them reported as not very well off or quite poor, which had a negative repercussion on or implications for their diet and quality of the food consumed, among others, that they may face these other health risks in the future is certainly high [6,11]. This prediction could be more accurate and precise through regression analyses of the response, which is not the case in this article. However, given the response and the multiple unhealthy and risky health behaviors and lifestyles reported by many of the respondents, chances are very high that many of the health risk factors could co-occur among many of them. It is commonly understood that negative health behaviors, attitudes, and lifestyles formed during childhood and adolescence have a profound impact on future health behaviors and lifestyles of the individuals. They will have significant negative impact on the individual health as well as familial economy and will be a big burden for the public policy to address in the future [3,16,18,33].

This study has some limitations. Firstly, the cross-sectional design could capture only a snapshot view of the study population and could not document changes in the variables over time. Given that conduct of student or youth health surveys are a rare phenomenon in Cambodia, findings from this study can provide some pioneering thoughts on the issue. Nevertheless, it must be underlined that many respondents reportedly adopted positive health behaviors and lifestyles, which may co-occur [17], although whether they co-occur was not examined in this study and warrant further analyses and research. Many of the health issues need to be examined into more details – examples include quality and adequacy of the food consumed, quality of sleep, and rigorousness of the physical activities. Other health and associated issues that may be addressed in the future research include mental health, family environment, and peer pressure. Secondly, the self-reported measures may have led to social desirability bias, particularly for sensitive information such as sexual behaviors and substance abuse. Future studies may want to triangulate the data through, for example, adoption of a mixed-method study. Third, the study may benefit from multiple regression analyses and multi-factorial association tests, which may shed better light on the results that would provide more nuanced and comprehensive findings. However, given that this paper attempts only to present broad basic description of the students’ health behaviors, this is beyond the scope of our study. Finally, this study included a sample of students selected from only two major universities located in the capital city and a province; therefore, findings from this study may not be generalized to a wider university student population in the country. However, because this is one of the few studies on Cambodian students’ health, it can provide policy implications for national health and education policy makers as well as the university community.

## Conclusions

Despite the above-mentioned limitations, our study provides preliminary insights into health-related behaviors among university students in Cambodia. We found that the prevalence of some health risks (such as risky sexual behaviors, tobacco use, alcohol consumption, and illicit drug use) among university students in this study was plausibly low. However, the rates of many other negative health behaviors (inactive lifestyle, high consumption of unhealthy food, and low consumption of healthy) were concerning. These health risks may predict later poor health including increased risks of non-communicable chronic diseases such as diabetes, hypertension, and other cardiovascular disease among this potential population. Further studies are needed to explore factors associated with specific health risk behaviors and effective intervention programs for promoting university student health in Cambodia. Effort may be needed to conduct such surveys on a more regular basis and on a larger scale to better inform public health programs and policies.

## Acknowledgements

The authors thank research assistants and the study participants for their contribution to this study. We also thank staff members at the Royal University of Phnom Penh and University of Battambang for their support during the data collection.

## Author Contributions

**Conceptualization:** Siyan Yi, Khuondyla Pal, Sovannary Tuot.

**Data curation:** Siyan Yi, Khuondyla Pal, Say Sok.

**Formal analysis:** Khuondyla Pal, Say Sok, Siyan Yi.

**Funding acquisition:** Sovannary Tuot, Siyan Yi.

**Investigation:** Siyan Yi, Khuondyla Pal, Pheak Chhoun, Rosa Yi, Sovannary Tuot, Say Sok.

**Methodology:** Siyan Yi, Khuondyla Pal, Pheak Chhoun, Rosa Yi, Say Sok, Sovannary Tuot.

**Project administration:** Khuondyla Pal, Sovannary Tuot, Pheak Chhoun, Rosa Yi.

**Resources:** Siyan Yi, Sovannary Tuot.

**Software:** Siyan Yi, Say Sok.

**Supervision:** Khuondyla Pal, Pheak Chhoun, Rosa Yi, Sovannary Tuot, Siyan Yi.

**Validation:** Siyan Yi, Say Sok, Sovannary Tuot.

**Visualization:** Sovannary Tuot, Say Sok, Siyan Yi.

**Writing – original draft:** Say Sok, Khuondyla Pal, Siyan Yi.

**Writing – review & editing:** Siyan Yi, Say Sok, Khuondyla Pal, Sovannary Tuot, Pheak Chhoun, Rosa Yi.

## Supporting information

**S1 File. STROBE checklist for cross-sectional studies**

## References

1. Department of Chronic Diseases and Health Promotion, World Health Organization. Preventing chronic diseases: a vital statement. Geneva: World Health Organization; 2005.

2. Mokdad AH, Marks JS, Stroup DF, Gerberding JL. Actual causes of death in the United States, 2000. JAMA. 2004;291(10):1238–45.

3. Yusuf S, Hawken S, Ounpuu S, Dans T, Avezum A, Lanas F, et al. Effect of potentially modifiable risk factors associated with myocardial infarction in 52 countries (the INTERHEART study): case-control study. Lancet. 2004;364(9438):937–52.

4. Danaei G, Vander Hoorn S, Lopez AD, Murray CJ, Ezzati M. Causes of cancer in the world: comparative risk assessment of nine behavioural and environmental risk factors. Lancet. 2005;366(9499):1784–93.

5. Lawlor DA, O’Callaghan MJ, Mamun AA, Williams GM, Bor W, Najman JM. Socioeconomic position, cognitive function, and clustering of cardiovascular risk factors in adolescence: findings from the Mater University Study of Pregnancy and its outcomes. Psychosomatic Medicine. 2005;67(6):862–8.

6. Peltzer K, Pengpid S. Heavy drinking and social and health factors in university students from 24 low, middle income and emerging economy countries. Community Ment Health J. 2016;52(2):239–44.

7. Pengpid S, Peltzer K, Kassean HK, Tsala JP, Sychareun V, Müller-Riemenschneider F. Physical inactivity and associated factors among university students in 23 low-, middle-and high-income countries. Int J Public Health. 2015;60(5):539–49.

8. Meng L, Maskarinec G, Lee J, Kolonel LN. Lifestyle factors and chronic diseases: application of a composite risk index. Prev Med. 1999;29(4):296–304.

9. Alamian A, Paradis G. Individual and social determinants of multiple chronic disease behavioral risk factors among youth. BMC Public Health. 2012;12:224.

10. World Health Organization. Global health risks: mortality and burden of disease attributable to selected major risks. Geneva: World Health Organization; 2009.

11. Peltzer K, Pengid S. Health behaviour interventions in developing countries. New York: Nova Publishers; 2011.

12. Rieger M. Risk aversion, time preference and health production: Theory and empirical evidence from Cambodia. Econ Hum Biol. 2015;17:1–15.

13. Fukuoka Y, Choi J, S Bender M, Gonzalez P, Arai S. Family history and body mass index predict perceived risks of diabetes and heart attack among community-dwelling Caucasian, Filipino, Korean, and Latino Americans–DiLH Survey. Diabetes Res Clin Pract. 2015;109(1):157–63.

14. Hawkins JD, Catalano RF, Miller JY. Risk and protective factors for alcohol and other drug problems in adolescence and early adulthood: implications for substance abuse prevention. Psychol Bull. 1992;112(1):64–105.

15. Perera S, Eisen R, Bawor M, Dennis B, de Souza R, Thabane L. Association between body mass index and suicidal behaviors: a systematic review protocol. Syst Rev. 2015;4:52.

16. Steptoe A, Wardle J, Cui W, Bellisle F, Zotti AM, Baranyai R, et al. Trends in smoking, diet, physical exercise, and attitudes toward health in European university students from 13 countries, 1990-2000. Prev Med. 2002;35(2):97–104.

17. Pengpid S, Peltzer K. Dietary health behaviour and beliefs among university students from 26 low, middle and high income countries. Asia Pac J Clin Nutr. 2015;24(4):744–52.

18. Steptoe A, Wardle J. Cognitive predictors of health behaviour in contrasting regions of Europe. Br J Clin Psychol. 1992;31:485–502.

19. Tapert SF, Brown GG, Kindermann SS, Cheung EH, Frank LR, Brown SA. fMRI measurement of brain dysfunction in alcohol-dependent young women. Alcohol Clin Exp Res. 2001;25(2):236–45.

20. Banta JE, Addison A, Job JS, Yel D, Kheam T, Singh PN. Patterns of alcohol and tobacco use in Cambodia. Asia Pac J Public Health. 2013;25(5 Suppl):33S–44S.

21. Schulenberg JE, Maggs JL. A developmental perspective on alcohol use and heavy drinking during adolescence and the transition to young adulthood. J Stud Alcohol Suppl. 2002;(14):54–70.

22. Douglas KA, Collins JL, Warren C, Kann L, Gold R, Clayton S, et al. Results from the 1995 National College Health Risk Behavior Survey. J Am Coll Health. 1997;46(2):55–66.

23. Nanakorn S, Osaka R, Chusilp K, Tsuda A, Maskasame S, Ratanasiri A. Gender differences in health-related practices among university students in northeast Thailand. Asia Pac J Public Health. 1999;11(1):10–5.

24. Peltzer K. Health behaviour in Black South African university students. S Afr J Psychol. 2000;30(4),46–49.

25. Yi S, Tuot S, Yung K, Kim S, Chhea C, Saphonn V. Factors associated with high-risk sexual behavior among unmarried male and female most-at-risk young people in Cambodia. Am J Public Health Res. 2014;2:211–20.

26. Yi S, Poudel KC, Yasuoka J, Palmer PH, Yi S, Jimba M. Role of risk and protective factors in risky sexual behavior among high school students in Cambodia. BMC Public Health. 2010;10:477.

27. Currie C, Hurrelmann K, Settertobulte W, Smith R, Todd J. Health and Health Behaviour among Young People. Copenhagen: World Health Organization; 2000.

28. National Institute for Health and Care Excellence, United Kingdom. Behaviour change: generviour change: general approaches. Manchester: National Institute for Health and Care Excellence; 2007.

29. Dinger MK, Waigandt A. Dietary intake and physical activity behaviors of male and female college students.. Am J Health Promot. 1997;11(5):360–2.

30. Steptoe A. An international study of personal heath behaviour, attitudes and well-being in university students. London: University of London; 2003.

31. Cambodia Development Resource Institute. Anatomy of Higher Education Governance in Cambodia. Phnom Penh: Cambodia Development Resource Institute; 2013.

32. Steptoe A, Wardle J. The European health and behaviour survey: the development of an international study in health psychology. Psychol Health. 1996;11:49–73.

33. Centers for Disease Control and Prevention. Youth Risk Behavior Surveillance: National College Health Risk Behavior Survey – United States, 1995. MMWR Morb Mortal Wkly Rep. 1997;46(SS-6):1–54.

34. Centers for Disease Control and Prevention. Global School-based Student Health Survey (GSHS). Atlanta: Centers for Disease Control and Prevention; 2014.

35. Craig CL, Marshall AL, Sjöström M, Bauman AE, Booth ML, Ainsworth BE, et al. International physical activity questionnaire: 12-Country reliability and validity. Med Sci Sports Exerc. 2003;35(8):1381–95.

36. Yi S, Poudel KC, Yasuoka J, Palmer PH, Yi S, Jimba M. Risk vs. protective factors for substance use among adolescents in Cambodia. J Subst Use. 2011;16(1):14–26.

37. Efroymson D, Ahmed S, Townsend J, Alam SM, Dey AR, Saha R, et al. Hungry for tobacco: an analysis of the economic impact of tobacco consumption on the poor in Bangladesh. Tob Control. 2001;10(3):212–7.

38. Sopheab H, Fylkesnes K, Vun MC, O’Farrell N. HIV-related risk behaviors in Cambodia and effects of mobility. J Acquir Immune Defic Syndr. 2006;41(1):81–6.

39. Gover PJ, Daan Aalders GJ. Does prevention have anything to do with it? Phnom Penh: Cambodian Communication Review; 2014.

40. Gover PJ, Daan Aalders GJ. Beyond Curiosity: A Re-examination of Positive Preventative Messages within the Cambodian Press. Phnom Penh: Cambodian Communication Review; 2016.

41. Agaku IT, King BA, Husten CG, Bunnell R, Ambrose BK, Hu SS, et al. Tobacco product use among adults--United States, 2012-2013. MMWR. Morbidity and mortality weekly report. 2014;63(25):542–7.

42. Singer M. Drugs and development: the global impact of drug use and trafficking on social and economic development. Int J Drug Policy. 2008;19(6):467–78.

43. Rehm J, Baliunas D, Borges GL, Graham K, Irving H, Kehoe T, et al. The relation between different dimensions of alcohol consumption and burden of disease: an overview. Addiction. 2010;105(5):817–43.

44. McCambridge J, McAlaney J, Rowe R. Adult consequences of late adolescent alcohol consumption: a systematic review of cohort studies. PLoS Med. 2011;8(2):e1000413.

45. Yahia N, Wang D, Rapley M, Dey R. Assessment of weight status, dietary habits and beliefs, physical activity, and nutritional knowledge among university students. Perspect Public Health. 2016;136(4):231–44.

46. World Health Organization. Global recommendations on physical activity for health. Geneva: World Health Organization; 2010.

47. Peltzer K, Pengpid S, Yung TK, Aounallah-Skhiri H, Rehman R. Comparison of health risk behavior, awareness, and health benefit beliefs of health science and non-health science students: An international study. Nurs Health Sci. 2016;18(2):180–7.

